# Genomic selection to optimize doubled haploid-based hybrid breeding in maize

**DOI:** 10.1101/2020.09.08.287672

**Authors:** Jinlong Li, Dehe Cheng, Shuwei Guo, Zhikai Yang, Ming Chen, Chen Chen, Yanyan Jiao, Wei Li, Chenxu Liu, Yu Zhong, Xiaolong Qi, Jinliang Yang, Shaojiang Chen

## Abstract

Crop improvement, as a long-term endeavor, requires continuous innovations in technique from multiple perspectives. Doubled haploid (DH) technology for pure inbred production, which shaves years off of the conventional selfing approach, has been widely used for breeding. However, the final success rate of *in vivo* maternal DH production is determined by four factors: haploids induction, haploids identification, chromosome doubling, and successful selfing of the fertile haploid plants to produce DH seeds. Traits in each of these steps, if they can be accurately predicted using genomic selection methods, will help adjust the DH production protocol and simplify the logistics and save costs. Here, a hybrid population (N=158) was generated based on an incomplete half diallel design using 27 elite inbred lines. These hybrids were induced to create F1-derived haploid families. The hybrid materials, as well as the 27 inbreds, the inbred-derived haploids (N=200), and the F1-derived haploids (N=5,000) were planted in the field to collect four DH-production traits, three yield-related traits, and three developmental traits. Quantitative genetics analysis suggested that in both diploids and haploid families, most of the developmental traits showed high heritability, while the DH-production and developmental traits exhibited intermediate levels of heritability. By employing different genomic selection models, our results showed that the prediction accuracy ranged from 0.52 to 0.59 for the DH-production traits, 0.50 to 0.68 for the yield-related traits, and 0.44 to 0.87 for the developmental traits. Further analysis using index selection achieved the highest prediction accuracy when considering both DH production efficiency and the agronomic trait performance. Furthermore, the long-term responses through simulation confirmed that index selection would increase the genetic gain for targeted agronomic traits while maintaining the DH production efficiency. Therefore, our study provides an optimization strategy to integrate GS technology for DH-based hybrid breeding.

## Introduction

Doubled haploid (DH) technology for homozygous inbred line production has been widely used in modern maize breeding because of the high efficiency and the low cost. Nevertheless, the molecular mechanisms by which the haploid is being induced remain mostly unclear. The maternal haploid induction rate is considered as a quantitative trait controlled by multiple genetic loci (Wu *et al*. 2014). Recently, major breakthroughs have been made by cloning the large effect QTLs for haploid induction, i.e., *qhir1* (Kelliher *et al*. 2017; Gilles *et al*. 2017; Liu *et al*. 2017) and *qhir8* (Zhong *et al*. 2019). These cloned genes and associated molecular evidence allowed researchers to re-evaluate the two competing hypotheses explaining the maternal haploid induction: (1) regular double fertilization followed by male chromosome elimination and (2) impaired double fertilization or single fertilization (Li *et al*. 2009a; Tian *et al*. 2018). It eventually led to a unified hypothesis that both fertilization defects and chromosome elimination could be involved in the maternal haploid induction (Chaikam *et al*. 2019b; Jacquier *et al*. 2020). However, more evidence is needed to elucidate the detailed molecular mechanisms for the phenomena.

To obtain the maternal DH lines *in vivo*, four essential steps are involved: (1) induction of maternal haploids by a male haploid inducer, (2) identification of haploid kernels or seedlings, (3) chromosome doubling of haploid seedlings (D0), and (4) selfing of fertile D0 plants to obtain DH seeds (Molenaar *et al*. 2019; Chaikam *et al*. 2019b). In the past several decades, continuous efforts have been made to improve the DH production from every perspective. To increase the haploid induction rate (HIR), several highly effective inducers have been developed, including MHI (Chalyk 1999), CAUHOI (Ming 2003), RWS (Röber *et al*. 2005), and PHI (Rotarenco *et al*. 2010), resulting in a dramatic increment of the HIR from 1-3% to 6-15% (Chaikam *et al*. 2019a). Besides the male factor, maternal germplasm also been confirmed to influence HIR (Kebede *et al*. 2011; Wu *et al*. 2014). After the induction, however, it is crucial to distinguish the induced haploids from the non-induced diploid kernels in order to make the following field trials more cost-effective. Currently, the widely used method for haploid identification is the *R1-nj* color marker system, where the haploid seeds show purple color on the aleurone only and the diploids exhibit purple color on both the aleurone and scutellum (Chaikam *et al*. 2015). To identify the haploid kernels more accurately and worthwhile, multiple different approaches or markers have been developed, including oil content (Ming 2003), Near-Infrared Spectroscopy (NIR) (Jones *et al*. 2012; Lin *et al*. 2019), and the double-fluorescence protein (DFP) marker (Dong *et al*. 2018). Chromosome doubling, or the haploid male fertility (HMF) and haploid female fertility (HFF), can be enhanced by chemical reagents, such as colchicine and herbicide (Saisingtong *et al*. 1996), but these chemicals are both harmful for human health and detrimental for the environment. Thus, spontaneous haploid genome doubling has been put on center stage (Ren *et al*. 2017; Boerman *et al*. 2020). Recent QTL studies suggested that the HMF is likely affected by several small effect loci (Ma *et al*. 2018; Ren *et al*. 2020). Relative to the low success rate of HMF, HFF exhibited a much higher success rate by pollinating from normal diploid plants (Geiger *et al*. 2006). And, therefore, HFF was not considered as a limiting factor for maternal DH production.

The ultimate goal of producing DH lines is to generate desired recombinants and to increase the genetic gain for traits of interest. However, because the genetic variation affects the success rates of maternal DH production for each step (Prigge and Melchinger 2012), it is necessary to select the appropriate DH-production methods based on the genotype without sacrificing the potential genetic gain. Genomic selection (GS), as a recently emerged technology to predict the performance of the plants without phenotyping, has been proved to be effective in plant breeding (Lin *et al*. 2016; Slater *et al*. 2016) and has the potential to increase the efficiency of DH-based selection. The widely used GBLUP model treats individual genotypes as random effects with their genomic relationship calculated from genome-wide markers (Henderson *et al*. 1984). Similarly, in the ridge regression BLUP (rrBLUP) model, markers were treated as random effects, with an assumption that each marker accounts for an equal amount of the genetic variance (Whittaker *et al*. 2000). The Bayesian alphabet models, i.e., Bayes A and Bayes B (Hayes *et al*. 2001), and Bayes C*π* (Habier *et al*. 2011), for genomic selection use hyperparameters to model marker variances differently (Kärkkäinen and Sillanpää 2012; Alves *et al*. 2019). Recently, some additional models, such as Neural Networks (Gianola *et al*. 2011), Bayesian LASSO (Gianola 2013), RKSH (Gianola *et al*. 2006), were developed and claimed to outperform the conventional models in some cases (Ogutu *et al*. 2012). In addition to enhance the statistical models, considering complex genetic effects can also have the potential to increase the prediction accuracy for GS. By considering dominance (Technow *et al*. 2012) and epistasis effects (Crossa *et al*. 2014), or even the genotype by environmental interactions (e Sousa *et al*. 2017), researchers improved the prediction performance for yield related and developmental traits in maize. Recently, Omics data started to be integrated into the genomic selection models. For example, transcriptomic and metabolomic data have been combined into genomic selection to boost the power of prediction (Hu *et al*. 2019). Additionally, by incorporating evolutionary information into the genomic selection model, the prediction accuracy has been improved for up to 4% for yield-related traits in maize (Yang *et al*. 2017). In this study, we sought to develop a strategy to integrate the GS for genetic improvement by considering the efficiency of DH production. With the empirical dataset collected from every step during DH production processes, results showed that DH-production traits could be accurately predicted using the GS models. To optimize the DH-based GS procedure, we constructed index traits and conducted index selection over 30 generations through simulation. The substantial long-term genetic gain using the index selection approach showed the feasibility of increasing multiple traits simultaneously. Our study streamlined a DH-based GS protocol that, if applied, has the great potential to facilitate plant breeding to meet the increasing food demands in the coming decades.

## Materials and Methods

### Plant materials and field experimental design

Here, 27 elite inbred lines were selected and crossed at Sanya (N18°37 E109°17) in 2017 Winter nursery according to an incomplete half diallel design to obtain N=158 hybrids (**Figure S1, Table S1**). In the following seasons, these hybrids and the inbred parents were grown in Beijing (N40°9 E116°23) during Summer 2018 and Sanya during Winter 2018. In the field, each accession was planted in a one-row plot with two replications; and 11 seeds were planted within each plot at a spacing of 60 cm between rows and 25 cm between plants. The CAU6, a haploid inducer with a high HIR (Zhong *et al*. 2019), was used to induce all hybrids and inbred parental lines at each location. After harvesting, the F1-derived and the inbred-derived haploid kernels were identified manually using the *R1-nj* color system (Chaikam *et al*. 2015). The identified haploid kernels were planted during Summer 2018 in Beijing and Winter 2018 in Sanya. For each F1-derived or inbred-derived haploid family, 48 individuals were planted in a three-row plot at a spacing of 60 cm between rows and 17 cm between plants. All of these haploids were pollinated by an inbred line C7-2, which has a high pollen count to ensure the success rate of pollination. The mature ears from the fertile plants were harvested manually.

### Phenotypic data collection

Along the four divided DH production processes, phenotypic data for ten different traits were collected. Briefly, data for three developmental traits, i.e., the plant height (PH), the ear height (EH), and the days to silking (DTS), were collected from the inbred parental lines and hybrids. To ensure the accuracy, three plants were measured for each row, and the average values were calculated for the following analyses. After crossing the hybrids and inbred lines with the inducer CAU6, mature ears were harvested to manually count the number of haploid kernels (*n*_*hk*_), diploid kernels (*n*_*dk*_), and embryo abortion kernels (*n*_*eak*_). With these counts, the maternal haploid induction rate (HIR) was calculated as 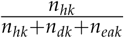. At each location, about 10 plants were induced for each hybrid. In the following seasons, putative haploid kernels were sowed, and diploid plants were identified using plant morphology observation method at around the V6-V8 stage (Ciampitti *et al*. 2011). With the field observation, the haploid plant rate (HPR) was calculated as 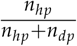, where *n*_*hp*_ was the number of haploid plants and *n*_*dp*_ was the number of diploid plants. The identified diploid plants were removed after data collection and the remaining haploid plants were pollinated by the elite inbred line C7-2. From the haploid plants, female and male fertility traits were collected. Briefly, the female fertility ratio (FFR) was calculated as 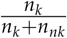, where *n*_*k*_ was the number of haploids contained more than one kernel, *n*_*nk*_ was the number of sterile haploids that didn’t generate any kernels. To evaluate haploid male fertility (HMF) trait, anther emergence ratio (AER) was computed using the formula of 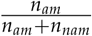, where *n*_*am*_ was the number of haploids that had observable anthers, *n*_*nam*_ was the number of haploids failed to detect any anthers. Finally, developmental traits were collected from the haploid plants, including PH, EH, and DTS; and yield-related traits were collected from the harvested mature ears, including the kernel row number (KRN) and the kernel number per row (KNPR). Because most of the ears didn’t have a full set of kernels, KRN and KNPR were evaluated using the embryo sac. Total kernel count (TKC) for each ear was simply computed by using KRN *×* KNPR.

### Phenotypic data analysis

The raw phenotypic data were analyzed using the linear mixed model with an R add-on package lme4 (Bates *et al*. 2015). Best linear unbiased predictors (BLUPs) were calculated for each F1-derived haploid family. In the model,

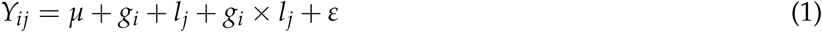

where, *Y*_*ij*_ is the mean phenotypic value of the *i*th F1-derived haploid family evaluated in the *j*th location; *µ* is the overall mean of the phenotypic trait in a F1-derived haploid family; *g*_*i*_ is the random effect of the *i*th F1-derived haploid family; *l*_*j*_ is the random effect of the *j* location; *g*_*i*_*×l*_*j*_ is the random interaction effect between the *i*th F1-derived haploid family and the *j*th location; and *ε* is the random error.

Similarly, BLUP values were calculated for each diploid genotype, where genotype, location, genotype and location interaction, and replication were treated as random effects. For traits collected in only one location, such as PH in inbred-derived haploid population, a simpler linear mixed model was employed, where genotype and plot were treated as random effects.

Heritability was calculated using variance component estimates from the above models. The following equation was used to estimate heritability on an individual plot basis,

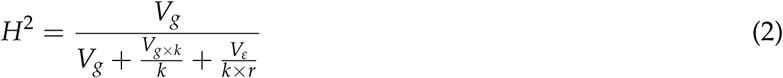

where *V*_*g*_ is the genotypic variance component, *V*_*ε*_ is the experimental error variance, *k* is the number of locations, *r* is the number of replications (*r* = 1 for the haploid family and *r* = 2 for the inbreds and F1 hybrids).

### Genotypic data processing and population structure analysis

Leaf tissues were sampled from the 27 inbred parental lines for DNA extraction using the CTAB method (Porebski *et al*. 1997). Then, genotyping was conducted using the Maize-60K SNP chip. SNPs with minor allele frequency (MAF) *<* 0.05 and per locus missing rate *>* 0.2 were filtered out using plink 1.90 (Chang *et al*. 2015). The cleaned SNP genotypes (*N* = 30, 887) were projected onto the F1 hybrids using a customized python package “impute4diallel” (https://github.com/jyanglab/impute4diallel).

By using the imputed SNPs, population structure analysis was conducted using the STRUCTURE software (Pritchard *et al*. 2000). In addition, Principal Components Analysis (PCA) was performed using the “princomp” function in R.

### Pedigree and genomic data enabled prediction models

The BLUP-based models were used to combine genomic or pedigree information into consideration (VanRaden 2008). In practice, the model is:

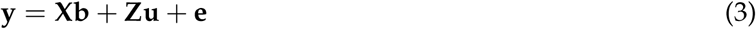

where **y** is a vector of the phenotype; **X** is a design matrix relating the fixed effects to each individual; **b** is a vector of fixed effects; **Z** is a design matrix allocating records to genetic values; **u** is a vector of genetic effects for each individual; and **e** is a vector of random normal deviates with variance *δ*^2^.

In our study, three different models based on above equation were applied, including genomic best linear unbiased prediction (GBLUP) with only additive effect (GBLUP-A), GBLUP with both additive effect and dominant effect (GBLUP-AD), and ridge regression best linear unbiased prediction (rrBLUP) (Endelman 2011). The models are,

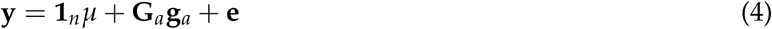

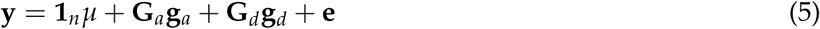

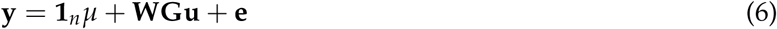

where **y** is the observed phenotypic value; **1**_*n*_ is a vector of ones (ignoring fixed effects); *µ* is the grand mean; **G**_*a*_ and **G**_*d*_ are the relationships matrices calculated by different methods; **W** is the design matrix associating accessions to observations; and **G** is the genotype matrix (row represents accessions and column represents biallelic SNP values); **g**_*a*_ and **g**_*d*_ are vectors of additive and dominance genetic effects, respectively; **u** denotes marker effects; and **e** is the residual error.

### The relationship matrix construction

The markers were coded as 1, 0, and *−*1, for SNP genotypes *A*_1_ *A*_1_, *A*_1_ *A*_2_, and *A*_2_ *A*_2_, respectively. The kinships of hybrids calculated from two genomic-based (**G**_*a*_ for additive, **G**_*d*_ for dominance) relationship matrices. The **G**_*a*_ and **G**_*d*_ were calculated as follows,

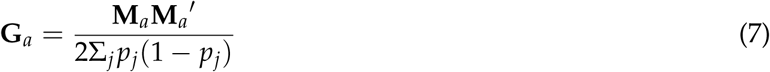

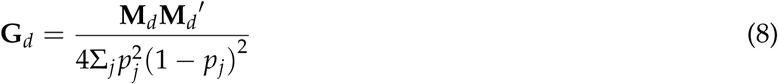

where *p*_*j*_ is the frequency of *A*_1_ allele at marker *j*; **M**_*a*_ and **M**_*d*_ are the *n × m* matrices (*n* is the number of individuals and *m* is the number of markers); **M** _*a*_ and **M**_*d*_ are the transposed matrices of **M**_*a*_ and **M**_*d*_. And the element of **M**_*a*_ or **M**_*d*_ for the *i*th individual at the *j*th marker is calculated as follows:

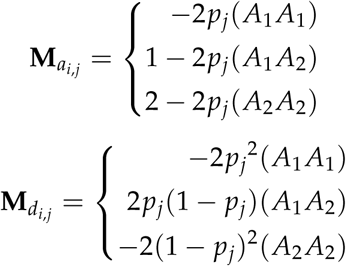

### Construction of the index traits

The indices were constructed using normalized values of the four DH-production traits and three yield-related traits using this formula for reasons that will be explained below.

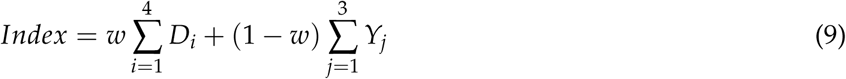

where *w* is the weighting parameter ranged from 0 to 1; in these experiments steps of 0.1 each were chosen. In the equation, *D*_*i*_ is the *i*_*th*_ DH-production trait (i.e., HIR, HPR, AER, and FFR); and *Y*_*j*_ is the *j*_*th*_ yield-related trait (i.e., TKC, KNPR, and KRN). Normalized trait values (with mean =0 and sd=1) were used to build the indices. The rrBLUP model was selected as the predictive model for the index traits. The 5-fold cross-validation method with 100 replications was used to assess the predictive ability for each index.

### Simulation for the DH-based long-term selection

To test the long-term responses of index selection, simulated selection experiments were conducted for 30 cycles. Our real-world hybrid population (N=158) was used as the initial training population. In the simulation, for each cycle, the top 20 hybrids based on the genomic estimated breeding values (GEBVs) were selected to be induced as haploids. The recombinations were simulated using an R add-on package “hypred” (Technow 2011) based on the published genetic map (Yu *et al*. 2008). The best recombinant haploid that survived for each hybrid-derived haploid family was doubled. These DH lines were crossed in a half-diallel manner to form the next cycle of hybrids. The GEBVs of simulated hybrids (N=190) were predicted using the rrBLUP model. Each simulation was repeated 20 times.

## Results

### The DH-production traits exhibited substantial genetic variation

In the first field experiment, 27 selected elite inbreds were crossed to generate F1 hybrids (N=158) based on a diallel design (**Figure 1** shows the experimental design and **Figure S1** specifies the hybrids made). These inbreds and hybrids were subsequently induced by CAU6 inducer (Zhong *et al*. 2019). The induced seeds were planted at two locations over three years. At each location, haploids were manually identified before flowering time and then pollinated by the C7-2 — an elite inbred line with a high pollen count (Li *et al*. 2009b). A number of phenotypic data were collected along with these processes, including four DH-production traits, three yield-related traits, and three plant developmental traits (**Table 1**, see **Materials and methods**). For these collected traits, the best linear unbiased predictors (BLUPs) were calculated for each genotype and haploid family (**Table S1**).

**Table 1.**
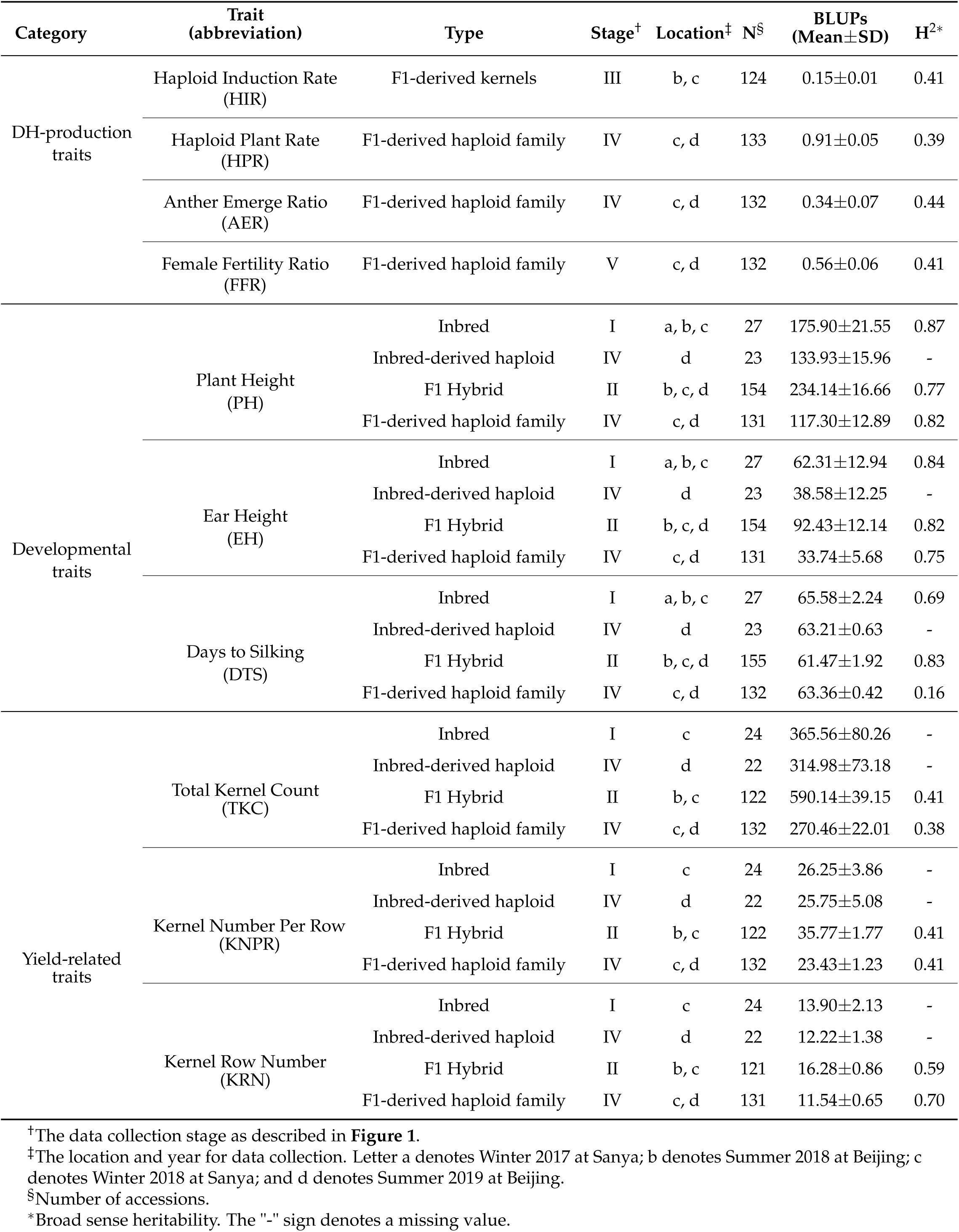
Summary of the phenotypic data analysis.

**Figure 1.**
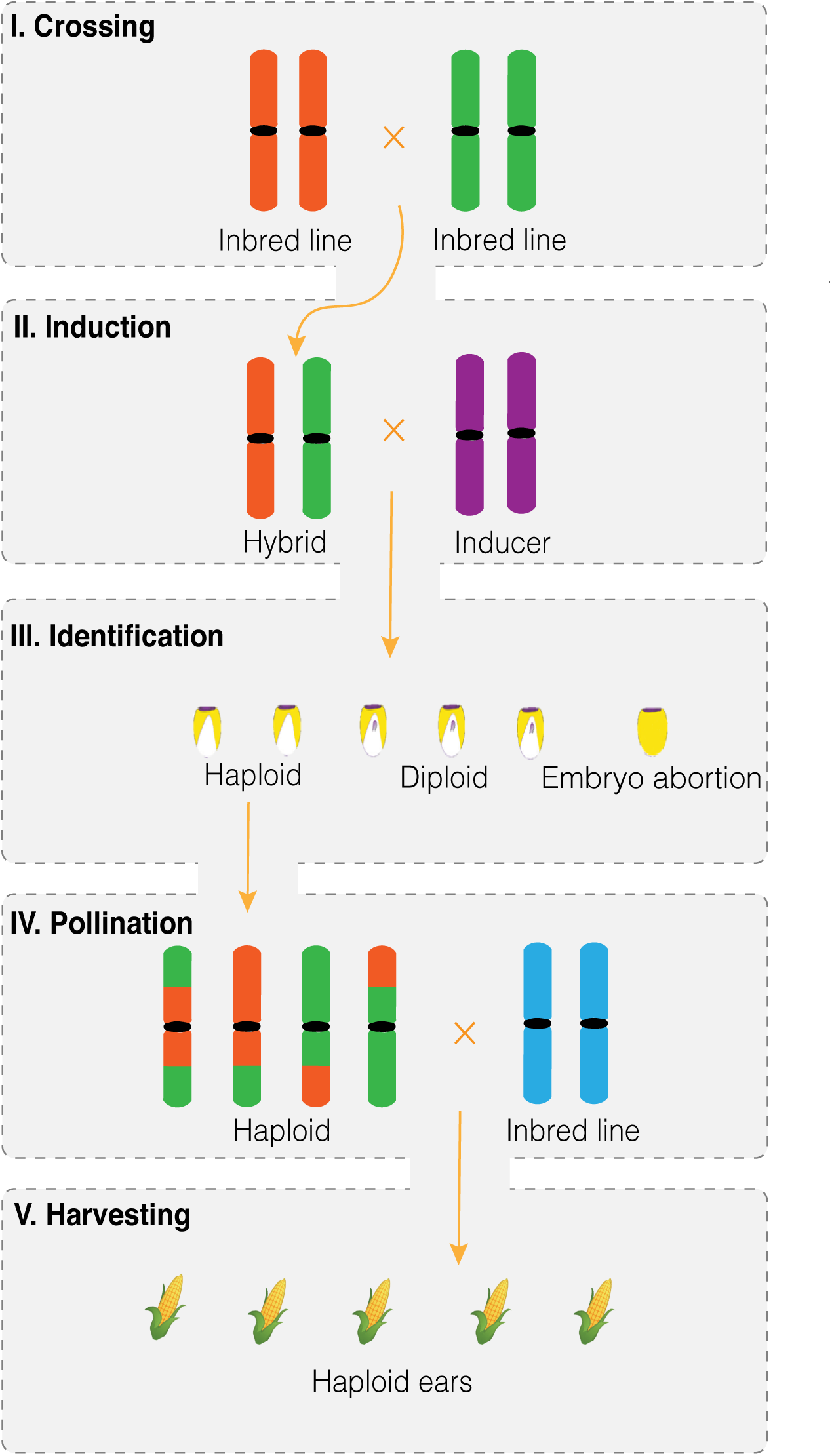
Schematic diagram of the experimental design. The diagram illustrates the five steps involved in the DH production, including crossing elite inbred lines to generate F1 hybrids (I), haploid induction using CAU6 as an inducer (II), haploid identification from harvested kernels (III), haploid pollination with an elite inbred line C7-2 (IV), and the harvesting of the mature ears from the haploid plants (V).

The BLUP values of the DH-production traits, including haploid induction rate (HIR), haploid plant rate (HPR), anther emergence ratio (AER), and female fertility ratio (FFR), exhibited bell-shaped distributions (**Figure 2**). The estimated mean HIR was 0.15±0.01, the lowest of the four DH-production traits, consistent with the previous observation that haploid induction was the step limiting trait for DH production (Prigge *et al*. 2012). In this experiment, the HPR, or the rate of the haploid plants out of the total plants, showed the highest value (mean = 0.91±0.05, ranged from 0.63 to 0.97), but substantial variations were observed. Two fertility traits, the male fertility trait (i.e., AER) and the female fertility trait (i.e., FFR), exhibited large variations with intermediate mean values of 0.34 ± 0.07 and 0.56 ± 0.06, respectively.

**Figure 2.**
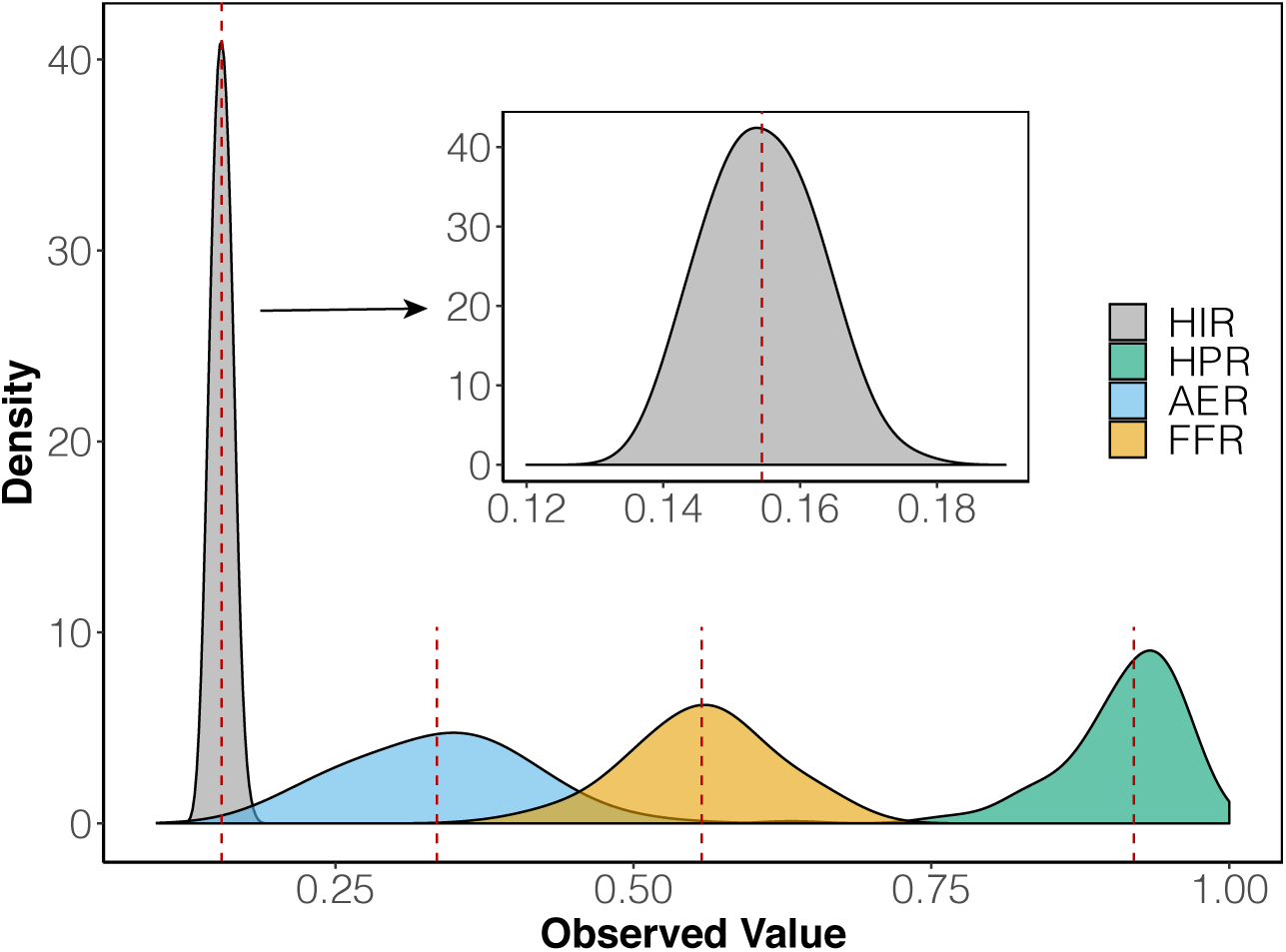
Phenotypic distributions of the four DH production-related traits. The probability density for values of the phenotypic trait of the F1-derived haploid families for the haploid induction rate (HIR), the haploid plant rate (HPR), the male fertility ratio (AER), and female fertility ratio (FFR) traits. The dotted red lines denote the mean values.

The developmental (i.e., PH, EH, and DTS) and yield-related traits (i.e., KRN, KNPR, and TKC) collected from the inbred parents, hybrids, inbred-derived haploids, and F1-derived haploids also exhibited distinct distributions (**Figure S2** and **Figure S3**), with diploids performing significantly better than haploids (Students’ t-test, P-value *<* 0.05), except for DTS, whereas the haploids were flowering late than hybrids but earlier than inbreds. Interestingly, inbred-derived haploids performed significantly better than hybrid-derived haploids for these developmental and yield-related traits, except for DTS (Student’s t-test, P-value *<* 0.01).

Additionally, correlation analysis suggested that the traits in diploids were correlated with traits in haploids derived from these diploids, especially for the developmental traits, whereas the Pearson correlation coefficients were above 0.6 for both PH and EH traits (**Figure S4**). For the yield-related traits (i.e., KRN, KNRP, and TKC), the correlations of BLUP values between haploids and diploids were weaker (*r* = 0.23*−* 0.51) but still statistically significant (Pearson correlation test, P-value *<* 0.05). Pair-wise correlations among the three categories of the traits showed that developmental and yield-related traits were positively correlated. However, in general, DH-production traits were negatively correlated with developmental traits and the yield-related traits, with some exceptions, for example, AER and DTS in hybrids (*r* = 0.30), AER and KRNP in hybrids (*r* = 0.24), AER and TKC in hybrids (*r* = 0.20), and FFR and KRN in haploids (*r* = 0.17) (**Figure S5**).

### The DH-production traits showed moderate levels of heritability

Phenotypes observed at multiple locations allowed us to estimate the heritability (see **Materials and methods**). The heritabilities of the four DH-production traits were 0.41, 0.39, 0.44, and 0.41 for HIR, HPR, AER, and FFR, respectively, largely consistent with previous studies (Wu *et al*. 2014, 2017; Ma *et al*. 2018) (**Table 1**). For the developmental and yield-related traits, heritabilities were estimated for both F1 hybrids and hybrid-derived haploid families. The yield-related traits, such as TKC and KNPR, exhibited intermediate levels of heritabilities (i.e., around 0.4 regardless of the populations), while the heritability for KRN was relatively higher (0.59 calculated from the hybrids and 0.70 from the F1-derived haploid families). For developmental traits, PH and EH exhibited high heritabilities in both haploid and diploid populations, ranging from 0.75 to 0.87. It was noticeable that heritability was extremely low for the DTS trait in the hybrid-derived haploid families, likely because male fertility after haploid induction was confounded with the flowering time trait. After excluding DTS, the heritability differences between F1 hybrids and hybrid-derived haploid families were insignificant (Paired t-test, P-value = 0.37).

### Genomic selection models for traits prediction

The parental inbred lines were genotyped using an SNP array (see **Materials and methods**). After SNP quality control, 30,887 remaining SNPs were projected onto the F1 hybrids. By using these projected SNPs, population structure analysis was conducted with STRUCTURE software (Pritchard *et al*. 2000). After testing group values (*k*) ranged from 2 to 10, *k* = 3 showed the highest likelihood, suggesting three subgroups within the F1 hybrid population (**Figure S6**), which was largely due to the crossing design (**Figure S1**). Principal component analysis (PCA) results also suggested three subgroups, with the first three principal components, explaining 20.76%, 20.12%, and 7.60% of the variances (**Figure S7**).

Next, we employed the projected SNP data to predict the phenotypic performance for the F1 hybrid and hybrid-derived haploid populations by taking account population structure and genetic relatedness into consideration. For the prediction, three models were used, including additive genomic BLUP (GBLUP-A) (VanRaden 2008), additive and dominance genomic BLUP (GBLUP-AD) (Da *et al*. 2014), and ridge regression BLUP (rrBLUP) (Whittaker *et al*. 2000) models (see **Materials and methods**).

Using a 5-fold cross-validation approach, the average prediction accuracies were 0.53± 0.04, 0.57± 0.05, 0.55± 0.03, and 0.52 ±0.04 for HIR, HPR, AER, and FFR, respectively (**Figure 3 (a)**). The prediction accuracy was significantly better than permutation results (Paired t-test, P-value < 0.01), suggesting that DH-production traits can be predicted accurately.

**Figure 3.**
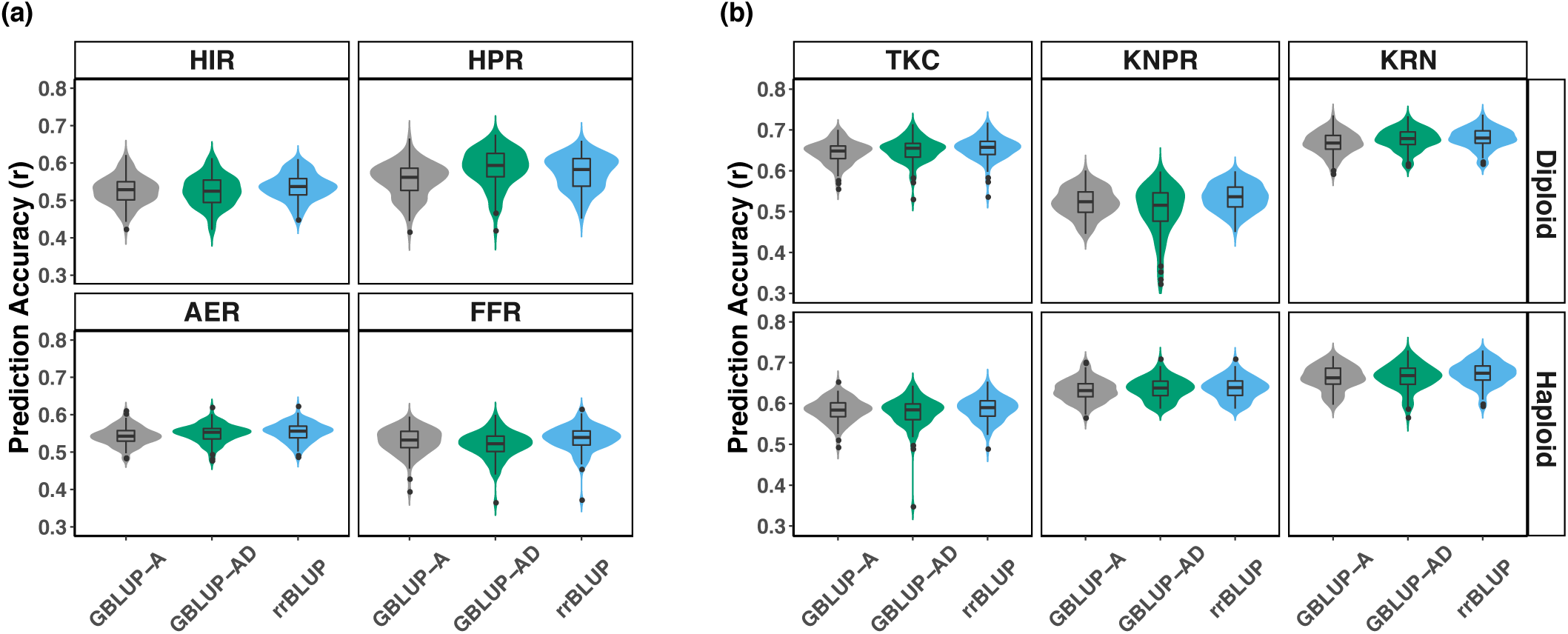
Predictive ability on DH-production traits and yield-related traits. **(a)** the prediction accuracy on the haploid induction rate (HIR), the haploid plant rate (HPR), the male fertility ratio (AER) and female fertility ratio (FFR) of the F1-derived haploid families. 3 models were used, genomic best linear unbiased prediction with additive effect (GBLUP-A), genomic best linear unbiased prediction with both additive and dominant effect (GBLUP-AD), ridge regression best linear unbiased prediction (rrBLUP); **(b)** the prediction accuracy on total kernel number (TKC), kernel number per row (KNPR), kernel row number (KRN) in diploid and haploid.

For the yield-related traits, GBLUP-AD outperformed the GBLUP-A model in both haploid and diploid populations, with the most considerable difference of 5.26% for the hybrid PH trait. These results were consistent with the assumption that dominance alleles affect these traits (Yang *et al*. 2017, 2018) (**Figure 3 (b)**). Similar patterns were also observed for predicting the developmental traits (**Figure S8**). The rrBLUP model, in most cases, performed equally well with the GBLUP-AD model. We therefore selected the rrBLUP model for the following analyses.

### The index traits integrated the DH production efficiency and agronomic performance

Genomic selection models can accurately predict the traits individually. However, given the DH-production traits were largely negatively correlated with the yield-related traits (**Figure S5**), promising recombinants with high yield performance may not be able to be produced through the DH pipeline. To increase the DH production efficiency without sacrificing the yield performance, the index traits were constructed by weighting both DH-production traits and yield-related traits (see **Materials and methods**). After changing the weighting coefficient (*w*) from 0 to 1 with step size of 0.1, rrBLUP results showed that prediction accuracy for the index trait peaked at *w* = 0.3, with the mean prediction accuracy = 0.71 (**Figure 4**).

**Figure 4.**
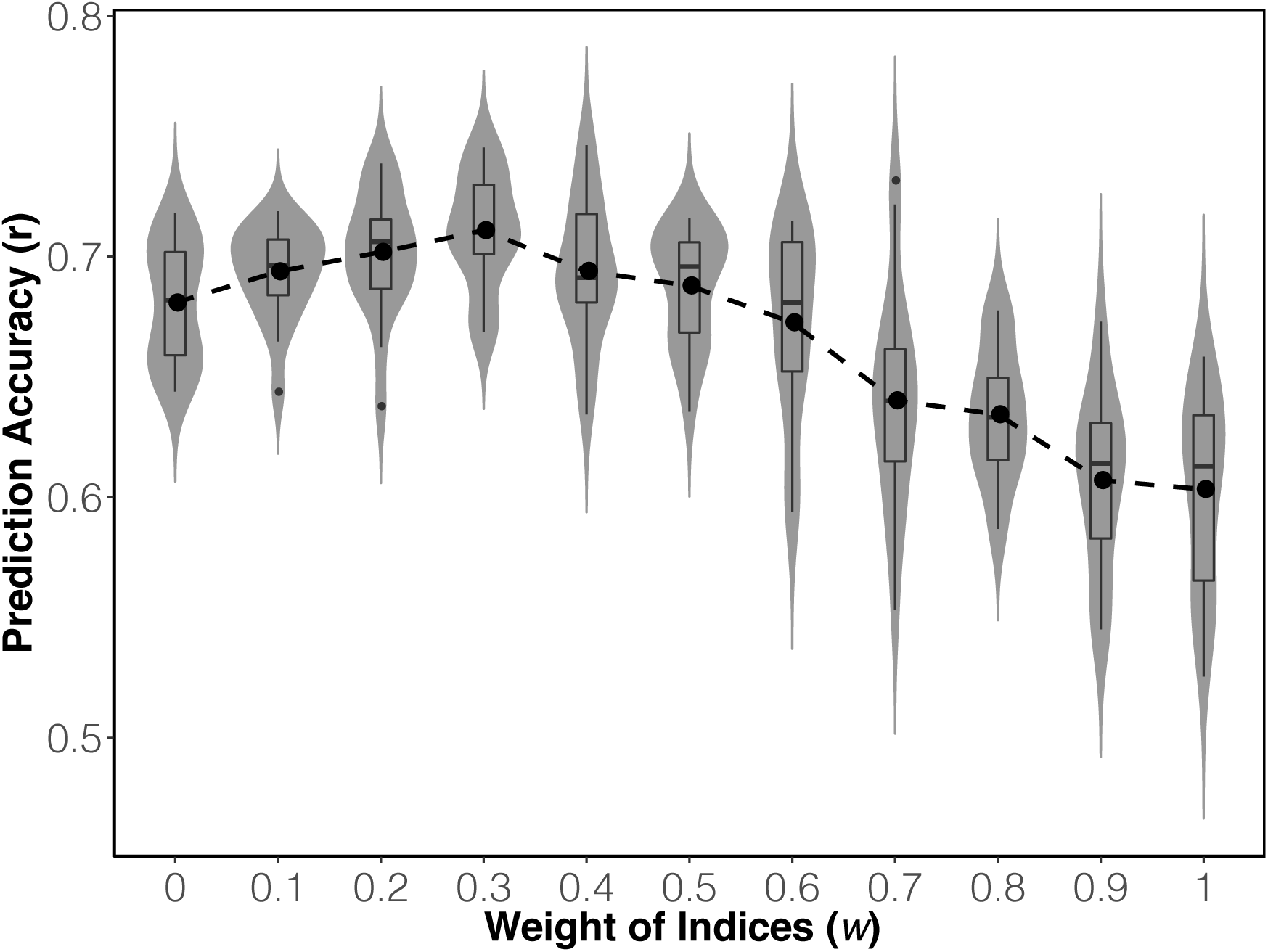
Predictive ability on indices. Horizontal axis denotes weight assigned to traits within the index.

In order to test the validity of index selection, we simulated the breeding selection process in 30 cycles (see **materials and methods**). In brief, a long-term selection experiment was simulated using our hybrid population as the initial training population (cycle 0). In the simulation, a fixed number of 1,000 seeds for each hybrid were induced per cycle. After considering the failure rate of each step during the DH-production process, the survived doubled-haploids were calculated for the genetic estimated breeding values (GEBVs) using the rrBLUP method. The top 20 DH lines were crossed based on a half diallel (*N* = 190) to advance to the next breeding cycle.

The results showed that after 30 cycles of simulation most of the traits reached to the plateau (**Figure 5**). Using the indices as the selection traits, regardless of the *w* value, GEBVs continued to increase, especially during the first several cycles of selection. When *w* is 0.5, where the DH-production traits and yield-related traits were equally weighted, GEBVs of the index traits were promoted to the highest value in each selection cycle, eventually reaching to 2.31 after 30 cycles of selection. If only one set of traits were selected, the longterm responses were comparatively low, i.e., at cycle 30, GEBVs = 2.03 when *w* = 0 (selecting only on the yield-related traits) and GEBV = 1.74 when *w* = 1 (selecting only on the DH-production traits).

**Figure 5.**
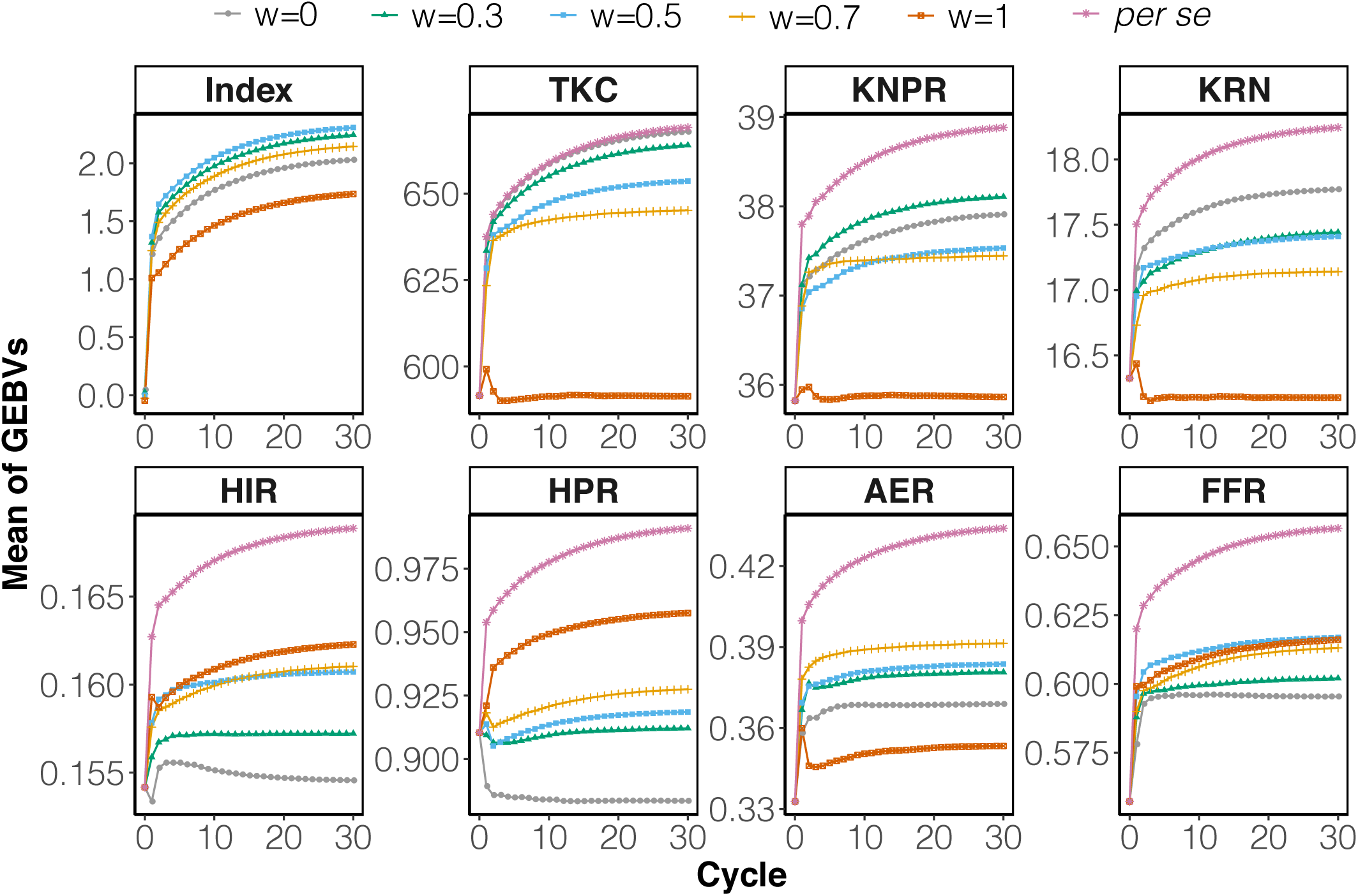
Genomic estimate breeding values (GEBVs) generated by selecting on different indices over 30 cycles. Five indices were tested, using weights set at 0, 0.3, 0.5, 0.7, and 1.0 (the trait was referred to by the weight chosen). By selecting index, GEBVs changes for diploid yield-related traits (i.e., TKC, KNPR, KRN) and DH production related traits (i.e., HIR, HPR, AER, FFR) were also calculated. *“Per se”* means select by the trait itself.

With the fixed number of induced plants (*N* = 1, 000 per hybrid), simulation results showed that the number of survived DH lines varied by the choices of *w*. The number was the highest at cycle 30 when *w* =0.7, increasing from 26 to 36 (a 38% improvement) (**Figure S9**). Alternatively, using the fixed number of DH lines produced per hybrid (*N* = 100), the index trait can be increased to 2.43 (*w* =0.5) after 30 generations of selection compared to 2.31 (*w* =0.5) using the fixed number of induced seeds, indicating that the production of more DH lines can improve the selection efficiency (**Figure S10**). However, the number of induced plants were almost tripled for each cycle, with *w* = 0.7 showing the most efficient DH production (**Figure S11**).

Not surprisingly, the traits achieved their highest values if selected on the traits *per se* rather than the indices. For TKC, one of the most important yield component traits, selection on trait *per se*, made almost no differences compared to selection on the *w* = 0 index trait. And the differences were minimum between selection on the trait *per se* and the *w* = 0.3 index trait, suggesting it is feasible to improve the yield component trait and the DH production efficiency simultaneously.

The long-term responses of individual traits vary by the choices of *w* values. For TKC and KRN, *w* = 0 made the greatest genetic gains, while for KNPR, *w* = 0.3 increased the most over 30 cycles of selection. Interestingly, *w* = 0.7 won the first 13 cycles of selection for KNPR; however, after cycle 13, *w* = 0.5 started to perform better. For DH-production traits, when *w* =1, HIR and HPR achieved the best results, increasing from 0.154 to 0.162 and 0.91 to 0.96 over 30 generations, respectively. The most effective *w* values for AER and FFR were 0.7 and 0.5, respectively. When *w* =1, due to negative correlations, the selection of the DH-production traits led to the negative responses for the TKC and KRN traits. Similarly, the negative response was observed for the HPR trait when selecting on the yield-related traits only (or *w* = 0). When the index coefficient of *w* was 0.3, 0.5, and 0.7, all traits were positively selected, suggesting that long-term selection using the index trait effectively increased both yield and DH production efficiency.

## Discussion

### Genomic selection technology could increase the long-term genetic gain

In the present study, DH lines were produced through a classical DH production pipeline for hybrid maize breeding (Prasanna *et al*. 2012). During the process, the DH-production traits, developmental traits, and yield-related traits were collected from haploids and diploids across multiple environments, which allowed us to calculate heritabilities for these traits. Results showed that four DH-production traits showed moderate levels of heritability (*H*^2^ ranged from 0.39 – 0.44), suggesting that DH production efficiency is under genetic control (Ma *et al*. 2018; Wu *et al*. 2014). For DH-production and yield-related traits, heritability estimations were highly consitent between haploids and diploids, suggesting limited contributions of allele interactions to the genetic variance.

The observation that some valuable hybrids were less efficient in generating DH lines provided an obstacle for further crop improvement through the recurrent selection approach (Bradshaw 2017). GS technology, a method to predict the phenotypic performance (i.e., the DH-production traits) without phenotyping, was proposed here to overcome the bottleneck. Promisingly, a moderate level of the prediction accuracy for each of the DH production traits was achieved (**Figure 3 (a)**), suggesting it is feasible to predict the DH-production traits before the haploid induction. Therefore, in practice, these predicted GEBVs can be leveraged to optimize the haploid induction, identification, doubling, and selfing processes. For example, more plants can be induced for hybrids with low inducing rates; the oil content approach can be used to improve haploid kernel identification instead of using the cost effective but less accurate coloring system (Ming 2003; Li *et al*. 2009a); and hybrids with low predicted chromosome doubling rates can be assisted with the chemical agent for chromosome doubling (Jumpatong *et al*. 1996). Even without using these additional enhancement approaches, simulation results showed that the long-term selection responses were significantly larger by allocating the appropriate number of inductions for each genotype than by inducing the same number of haploids for all genetic backgrounds. The improvement for the total kernel count after 30 cycles of simulated selection was up to by 13%, a substantial improvement achieved just by allocating resources differently.

### Index selection improved multiple traits simultaneously

The ultimate goal of DH-based plant breeding is to increase agronomic traits performance. In practice, however, low DH-production efficiency creates the logistics burden. To improve both types of traits, index traits considering these two were constructed. The cross-validation results suggested that the weighting coefficient (*w* = 0.3) provided the best prediction accuracy for the index trait, – 4.41% and 18.33% improvements compared to selecting only on yield-related traits (*w* = 0) and DH-production traits (*w* = 1), respectively. However, the long-term responses for the index traits performed the best when weighing both types of traits equally (*w* = 0.5). According to Falconer and Mackay (Falconer and Mackay 1996), the long-term response to selection is influenced by the intensity of selection, the heritability of the trait, and the standard deviation of the breeding value. Therefore, it is reasonable that higher prediction accuracy alone can’t guarantee the best long-term response.

For the individual trait of interest, index selections (i.e., *w* = 0.3, 0.5, or 0.7 in our simulations) led to multiple traits improvement simultaneously, although the magnitudes of responses were smaller than directly selected on the trait *per se* (Su *et al*. 2012; Cui *et al*. 2020). Index selection, however, will avoid the situations of traits declining if they were negatively correlated with the trait that was under direct selection.

### Genomic selection models performed equally well in predicting DH-production traits

The predictive ability of a given model can be affected by heritability, training population size, the density of the markers, and the mating design alongside with the genetic architecture of the trait (Jumpatong *et al*. 1996). Previous studies and our own data showed that DH-production traits were complex traits controlled by many small-effect QTLs with relatively low heritability (Boerman *et al*. 2020; Ren *et al*. 2020). After comparing multiple GS models, our results suggested rrBLUP and GBLUP (including GBLUP-A and GBLUP-AD) only exhibited subtle differences in predicting the DH-production traits. Overall, the rrBLUP was considered a stable model, because in most cases, it performed the best or close to the best performing models. For the developmental and yield-related traits, dominance GBLUP (GBLUP-AD) exhibited higher prediction accuracies than the additive GBLUP (GBLUP-A) model in the diploid populations.

Overall, this study provided evidence that it is feasible to use GS technology to optimize the DH-based plant breeding. If implemented appropriately, the long-term genetic gain can be substantial, as illustrated by the simulations. This overall strategy can be applied,not only for maize but for other crop species, to breed the next generation of crop species faster and more cost-effectively.

## ACKNOWLEDGEMENTS

This research was funded by the National Key Research and Development Plan (2016YFD0101200), the Modern Maize Industry Technology System (CARS–02–04), and the Beijing Agricultural Reform and Development Special Transfer Payment Fund from Beijing Municipal Bureau of Agriculture and Rural Affairs to S.C.. This project was also partly supported by an Agriculture and Food Research Initiative Grant (Number 2019-67013-29167) from the USDA National Institute of Food and Agriculture, and by the University of Nebraska-Lincoln start-up fund to J.Y.. The computational work was completed utilizing the Holland Computing Center of the University of Nebraska, which receives support from the Nebraska Research Initiative. We thank Beijing Tongzhou International Seed Science and Technology Co., Ltd. for the genotyping effort.

## AUTHOR CONTRIBUTIONS

S.C., J.Y. and J.L. designed this work. J.L., D.C., S.G., M.C., C.C., Y.J., W.L, Y.Z. and X.Q. generated the data. J.L., J.Y., and Z.Y. analyzed the data. S.C. and C.L. provided conceptual advice. J.Y., J.L., and S.C. wrote the manuscript.

## DATA AVAILABILITY

The data and code of this project were released at the GitHub repository (https://github.com/lijinlong1991/DH-production-GS).

## ABBREVIATIONS

AER: anther emergence ratio
BLUPs: best linear unbiased predictors
CTAB: cetyltrimethylammonium bromide
DFP: double-fluorescence protein
DH: doubled haploid
DTS: days to silking
EH: ear height
FFR: female fertility ratio
GBLUP: genomic best linear unbiased prediction
GBLUP-A: genomic best linear unbiased prediction with only additive effect
GBLUP-AD: genomic best linear unbiased prediction with both additive effect and dominant effect
GEBV: genomic estimated breeding value
GS: genomic selection
HFF: haploid female fertility
HIR: haploid induction rate
HMF: haploid male fertility
HPR: haploid plant rate
KRN: kernel row number
KNPR: kernel number per row
MAF: minor allelefrequency
PCA: principal component analysis
PH: plant height
rrBLUP: ridge regression best linear unbiased prediction
SNP: single nucleotide polymorphism
TKC: total kernel count

